# Micro to macro scale anatomical analysis of the human hippocampal arteries with synchrotron Hierarchical Phase-Contrast Tomography

**DOI:** 10.1101/2024.08.06.605570

**Authors:** A. Bellier, P. Tafforeau, A. Bouziane, T. Angelloz-Nicoud, P.D. Lee, C. Walsh

## Abstract

**Purpose:** To date, no non-invasive imaging modality has been employed to profile the structural intricacies of the hippocampal arterial microvasculature in humans. We hypothesised that synchrotron-based imaging of the human hippocampus would enable precise characterisation of the arterial microvasculature.

**Methods:** Two preserved human brains from, a 69-year-old female and a 63-year-old male body donors were imaged using hierarchical phase-contrast tomography (HiP-CT) with synchrotron radiation at multiple voxel resolutions from 25.08 μm down to 2.45 μm. Subsequent manual and semi-automatic artery segmentation were performed followed by morphometric analyses. These data were compared to published data from alternative methodologies.

**Results:** HiP-CT made it possible to segment in context the arterial architecture of the human hippocampus. Our analysis identified anterior, medial and posterior hippocampal arteries arising from the P2 segment of the posterior cerebral artery on the image slices. We mapped arterial branches with external diameters greater than 50 μm in the hippocampal region. We visualised vascular asymmetry and quantified arterial structures with diameters as small as 7 μm.

**Conclusions:** Through the application of HiP-CT, we have provided the first imaging visualisation and quantification of the arterial system of the human hippocampus at high resolution in the context of whole brain imaging. Our results bridge the gap between anatomical and histological scales.

## Introduction

The hippocampus, located in the temporal region of the cerebrum’s limbic lobe, plays a pivotal role in memory and learning. This C-shaped structure, situated medially within the temporal lobe, consists of two intertwined allocortical layers: the *gyrus dentatus* and the *cornu ammonis* (CA1 to CA4). Typically, it is segmented into three parts: the head, body, and tail, each having intraventricular and extraventricular components [20]. The *cornu ammonis* is further divided into four fields based on their cellular morphology and susceptibility to hypoxia [23]. The hippocampus, boasts extensive connections with regions such as the entorhinal area, diencephalon, and brainstem. Notably, it connects with the mammillary body, anterior thalamic nuclei, hypothalamus, amygdala, and the contralateral hippocampus via the fimbria [3]. Afferences to the hippocampus include those from the cortical area, amygdaloid complex, medial septal region, thalamus, and more. Half of the septal nuclei inputs are cholinergic [15] and the other half are GABAergic [9]. Many of these connections, channelled through the trisynaptic circuit [14], facilitate the hippocampus’s role in functions like scene recognition [21], episodic and long-term memory [5], and creating a cognitive map [12]. This structure is pivotal for storing memories related to social interactions and hierarchies [17].

The hippocampus primarily receives vascularisation from the posterior cerebral artery (PCA) and also from the anterior choroidal artery (A-Ch-A). The PCA branches into the inferior temporal, posterolateral choroidal, and splenial arteries, which further give rise to the superficial hippocampal arteries. Additionally, a direct superficial hippocampal artery arises from the A-Ch-A [4]. The arterial groups from PCA and A-Ch-A follow each other on the surface of the subiculum, they anastomose and send perforating arteries to vascularise the hippocampus’s body and tail while the anterior hippocampal arteries, vascularise uncal apex, entorhinal area and anastomose with uncal branches of the A-Ch-A. Finally, the superficial arteries lead to intra-parenchymal arteries that can be divided into 4 groups: large ventral intrahippocampal arteries (for CA1 and CA2), small ventral intrahippocampal arteries (for the proximal part of the gyrus dentatus), large dorsal intrahippocampal arteries (for CA4, CA3 and the distal part of the gyrus dentatus), and small dorsal intrahippocampal arteries (for CA3 and CA4 via the fimbriodentate sulcus) [4]. Due to its hypoxic vulnerability [18], the hippocampus is implicated in various disorders [3]. Enhanced vascularisation, notably from A-Ch-A and PCA, is associated with better cognitive performance [7, 18] and hippocampal volume. Dysregulation in CA1’s glutamatergic neurons is linked to schizophrenia [11]. Other studies have identified left hippocampal atrophy in Parkinson’s-related cognitive impairment [16] and explored the hippocampus as a target for deep brain stimulation in epilepsy [8, 22].

Vascularisation studies of the hippocampus have traditionally employed conservative imaging techniques, including conventional computed tomography (CT) and magnetic resonance imaging (MRI), with limited resolution. For superior resolution, methods, such as light microscopy, electron microscopy, and injections-corrosion combined with scanning electron microscopy, have been utilised, but these methods do not preserve the sample. Recently, scientists have advanced non-destructive imaging techniques that offer enhanced visualisation from macro to micro scale using synchrotron radiation: Hierarchical Phase-Contrast Tomography (HiP-CT). We hypothesised that synchrotron-based imaging of the human hippocampus would enable more precise assessment of the hippocampal vasculature in the context of the complete brain imaging. The objective of the study was to verify this hypothesis.

## Materials and Methods

### Study design

We performed a descriptive anatomical study. The brains used were obtained from body donation to the Department of Anatomy (LADAF) of the Grenoble Alpes University. The methodological quality of this observational cadaveric study was assessed by Quality Appraisal for Cadaveric Studies (QUACS) [25].

### Sample preparation

A preservation process was initiated within 36 hours post-mortem, utilising a solution of formaldehyde and lanolin, administered via the carotid arteries. Subsequent anatomical dissection allowed for a cranial flap craniotomy and extraction of the entire brain. This specimen was then submerged in a 4% buffered formaldehyde solution, with the solution’s volume being at least four times that of the brain, and stored in a refrigerated chamber. Following a minimum fixation period of three days, the brain was prepared following the protocol previously presented [2, 24]. The brain was placed in a series of 4 successive ethanol bathes starting with 50% with increasing concentration up to 70% and degassed at each bath. To prevent imaging artefacts from movement, the brain was stabilised using agar gel. The whole mounted organ with the agar in the 70% ethanol was degassed before definitive sealing of the container.

### Scanning, reconstruction and segmentation

Imaging was performed on the BM05 beamline or on the BM18 beamline at the European Synchrotron Radiation Facility (ESRF) following the HiP-CT protocol [2, 24]. This new technique has been made possible thanks to the first high-energy, 4th generation synchrotron upgrade at ESRF, named the Extremely Brilliant Source. Initially, a whole brain was imaged at 25.08 μm^3^/voxel (edge length) on BM05 (data previously published in the Human Organ Atlas, DOI 10.15151/ESRF-DC-572252655) with a propagation distance of 3.5 m (brain 1). The right hippocampus region of the same brain was also imaged at 6.05 μm^3^/voxel and 2.45 μm^3^/voxel. A second brain (brain 2) was imaged at 23.42 μm^3^/voxel on BM18 with a propagation distance of 20 m, making all the small structures much more visible than with the BM05 setup (Figure 1). Tomographic reconstruction was performed using single distance phase-retrieval coupled with filtered back-projection algorithm as described in [24].

**Fig. 1.**
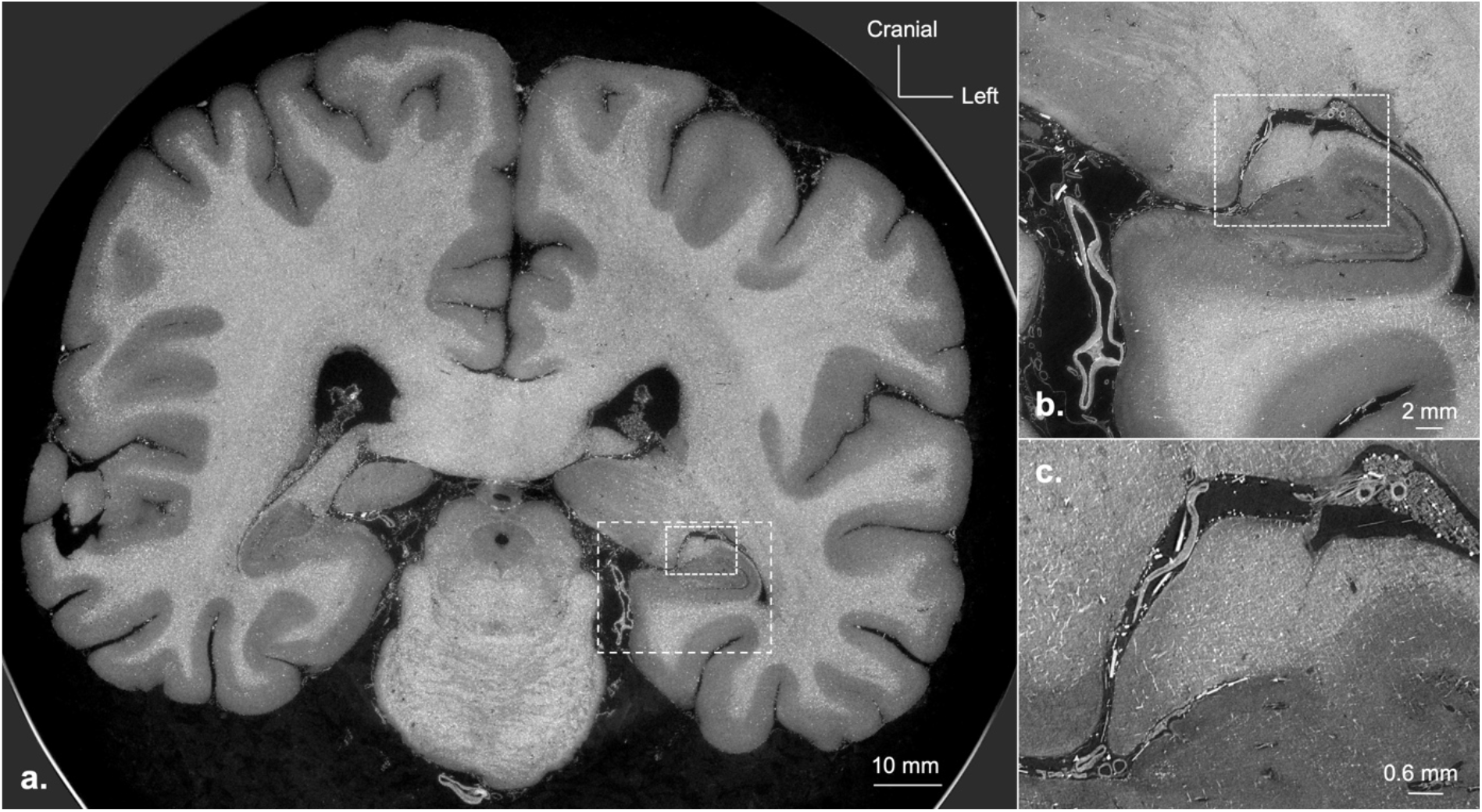
Frontal sections of human brain (second specimen) with hierarchical phrase-contrast tomography at 23.42 μm resolution **a**. Whole-brain; **b**. First zoom on hippocampus area; **c**. Second zoom on hippocampus area

For the brain 1, three-dimensional reconstruction of the arteries was obtained by manual virtual segmentation using ITK-SNAP Software (version 3.6.0) on a WACOM Cintiq Pro 24’’ graphic tablet using an interactive stylet allowing for more precise segmentation. A trained author-operator performed the manual segmentation. It consisted of a manual contouring of the anatomical structures identified on axial sections. The arteries were identified, thanks to the different tunics composing their walls, up to the intra parenchymal level. The segmentation was double checked by an experienced anatomist. The external diameter of the hippocampal arteries was assessed using the ImageJ Software on 10 axial measures with manual segmentation at their origin (i.e., after their branching to the higher generation arteries). For the brain 2, semi-automatic segmentation by local region growing was used, performed by an experienced anatomist using VG Studio Max software version 2023.3. The measurements were carried out on this software using the same method as described above.

We mainly used descriptive statistics including means with a standard deviation for quantitative variables. All the calculations and graphs were done with the RStudio software (Version 1.3.959© 2009-2020 RStudio, PBC).

## Results

### Study sample

The study included two brains. Brain 1 corresponded to that of a 69-year-old female (LADAF 2020-31) with type 2 diabetes, cystectomy, right colectomy, and peritoneal carcinoma treated by omentectomy and radiation. The body measured 145 cm and weighed 40 kg. Brain 2 corresponded to that of a 63-year-old male (LADAF 2021-17) with non-metastatic pancreatic cancer. The body measured 178 cm and weighed 60 kg.

### Image analysis

Utilising the HiP-CT technique, we achieved high-resolution slices with a contrast that readily differentiates the grey matter, the white matter, and the entirety of the vasculature. Based on imaging analysis, we were able to visualise the two perimesencephalic segments of PCA (Figure 2): precommunicating segment (P1) in the intercrural cistern, between origin of PCA and the meeting of posterior communicating artery, and postcommunicating segment (P2) which is located in the ambient cisterns. In its P2 segment, the PCA leads to the posterolateral choroidal artery, the splenial artery and the inferior temporal arteries (ITA), which are usually divided into the anterior, medial, and posterior branches. In addition to its uncal branch, the anterior choroidal artery, originating from the posterior side of the internal carotid artery, also reaches the choroids plexuses of the temporal horn of the lateral ventricle by following the choroid fissure. In the ambient cistern, PCA circulate along the parahippocampal gyrus and terminates in P3 segment of PCA, corresponded to the quadrigeminal segment.

**Fig. 2.**
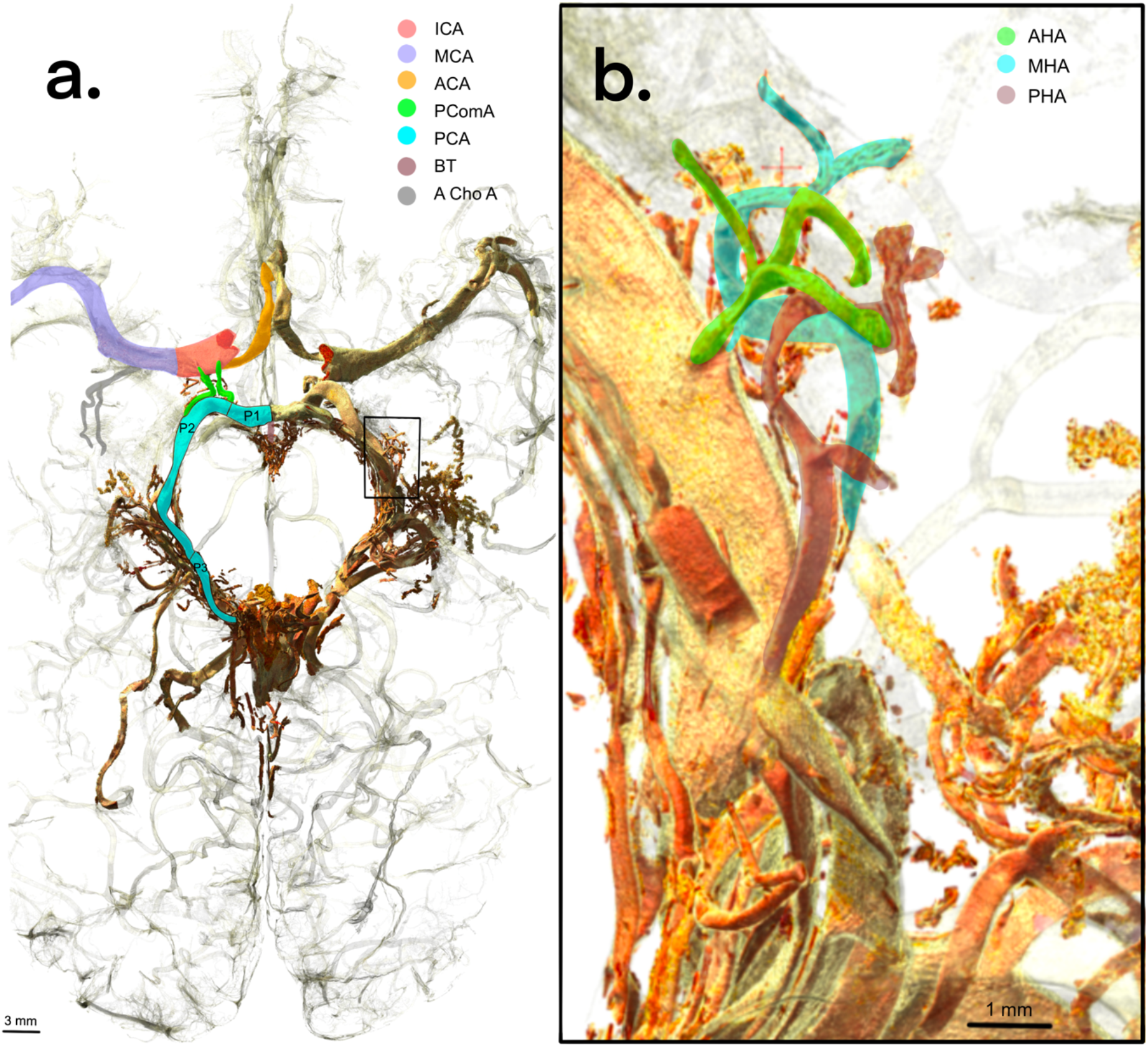
**A**. Three-dimensional segmentations of the human cerebral arterial tree (cranial view) in our specimens. **a**. Whole-brain; **b**. Zoom on right hippocampus arterial tree *ICA: internal carotid artery; MCA: middle cerebral artery; ACA: anterior cerebral artery; PComA: posterior communicating artery; PCA: posterior cerebral artery; BT: basilar trunk; A Cho A: Anterior choroidal artery; AHA: anterior hippocampal artery; MHA: middle hippocampal artery; PHA: posterior hippocampal artery*

### Vascular organisation

We have identified the three usual groups of arteries that vascularise the hippocampus: the anterior, the medial and the posterior hippocampal superficial arteries. In the right hemisphere of both subjects, the anterior hippocampal artery originated from the anterior ITA, the medial hippocampal artery originated from the middle ITA and the posterior hippocampal artery emanated directly from the trunk of PCA. The posterior hippocampal artery was identified as two branches, one emanating from the splenial artery and the other directly from the PCA (Figure 3). In the left hemisphere, the territory of anterior hippocampal artery is vascularised by a branch originating directly from the trunk PCA and a second branch coming from the anterior ITA. The medial hippocampal artery arises from the medial ITA. For the posterior hippocampal artery, we have identified the same two branches as for the right hemisphere: one emanating from the splenial artery and the other directly from the PCA. This anatomical arrangement was relatively similar in both brains studied (Figure 3).

**Fig. 3.**
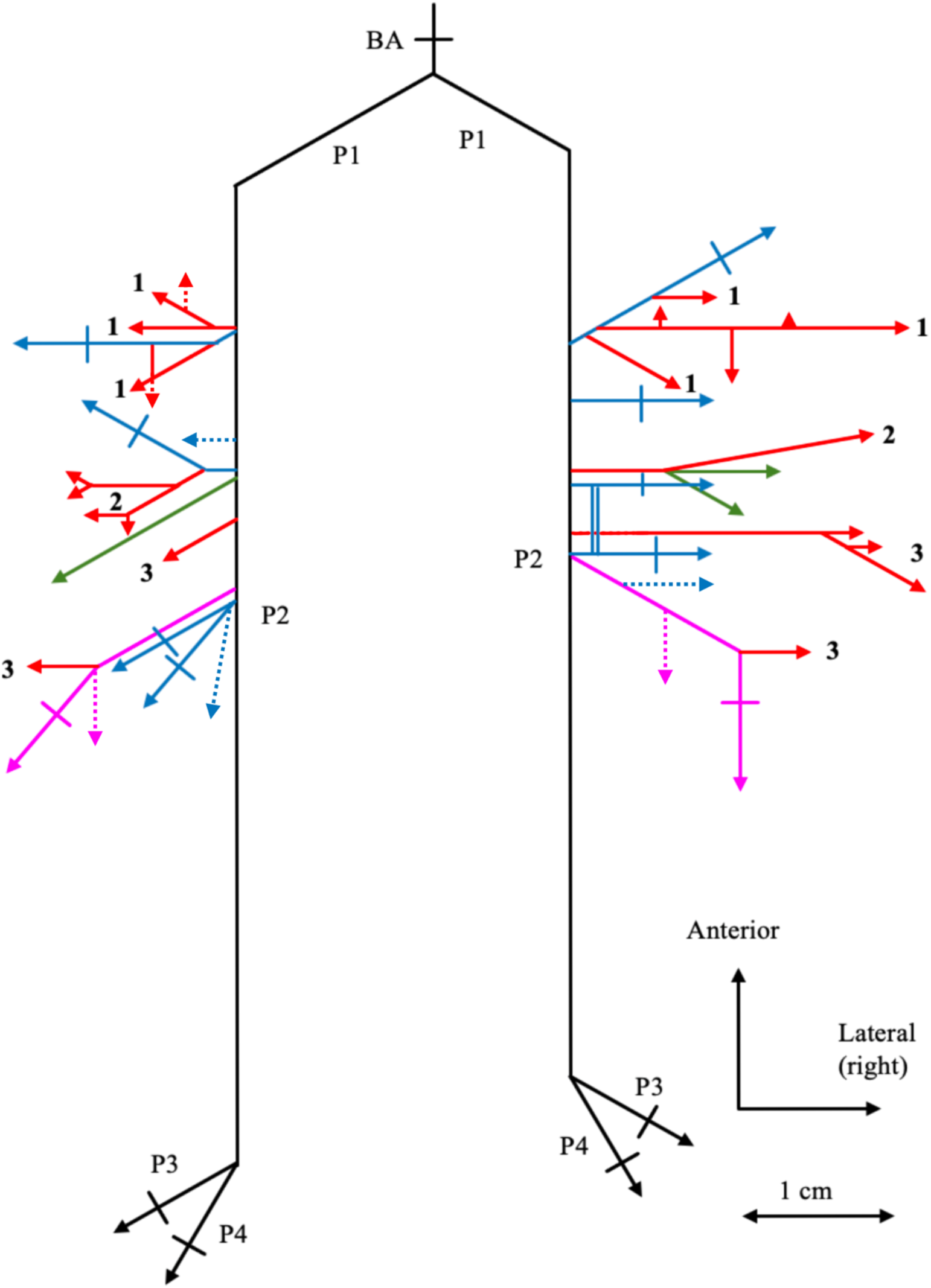
Diagram of the human hippocampal arterial tree in our specimens 1: anterior hippocampal artery; 2: medial hippocampal artery; 3: posterior hippocampal artery; BA: basilar artery; P1, P2, P3 and P4: segments of the posterior cerebral artery *Colours: Blue= inferior temporal artery; Green= choroidal plexuses artery; Pink= splenial artery; Red = hippocampal artery* *Solid line= average configuration when present in both specimens* *Dotted line= present in a single specimen*

### Quantitative analysis

In Table 1, the mean diameter (measure unit μm) of the main hippocampal arteries for both hemispheres is presented. The medial hippocampal arteries were the smallest with a mean of 129.6 µm (SD: 0.1) for both hemispheres, while the posterior hippocampal arteries were the largest with a mean of 232.2 µm (SD: 0.1) for both hemispheres. In addition, the thinnest arteries diameter has been measured at 7 μm on right hippocampus on the scans at 2.45 μm voxel size (Figure 2).

**Table 1.** Quantitative analysis of arterial vessels involved in hippocampal vascularisation. *AHA: anterior hippocampal artery; MHA: medial hippocampal artery; PCA: posterior cerebral artery; PHA: posterior hippocampal artery; SD: standard deviation*

| Hemisphere | Artery | Mean (SD) diameter ($\mu\text{m}$) |
| --- | --- | --- |
| Right | P2 segment (PCA) | 848.3 (0.2) |
|  | AHA | 155.8 (0.1) |
|  | MHA | 149.8 (0.1) |
|  | PHA | 237.6 (0.1) |
| Left | P2 segment (PCA) | 930.1 (0.5) |
|  | AHA | 131.1 (0.1) |
|  | MHA | 109.4 (0.1) |
|  | PHA | 226.7 (0.1) |

## Discussion

Utilising HiP-CT, the vascular architecture of the human hippocampus in the context of a whole brain imaging was imaged at a voxel resolution of 25.08 μm with 3.5 m of propagation on BM05 and 23.42 μm with 20 m of propagation on BM18, signifying a voxel resolution enhancement by a factor of 20 when compared with standard clinical CT scanners. The longer propagation distance on BM18 significantly increased the visibility of small structures, especially relevant for fine vasculature 3D segmentation. HiP-CT imaging yielded unprecedented resolution, enabling visualisation of intraparenchymal vessels within their native tissue context. Compared to laboratory micro-CT with contrast agent, the technique used in our study enables us to scan a large volume and therefore an entire brain. Such advances position synchrotron HiP-CT comparably to high-resolution methods like histology, contrast agent injection techniques and corrosion casts; however, it is unique in its non-destructive nature and ability to maintain relative anatomical integrity, offering 3D structural insights into vessels currently down to 10 µm in diameter [4, 13]. This marked the first instance where a three-dimensional visualisation of the human hippocampal arteries and the hippocampus itself has been achieved at this resolution. 3D rotational angiography techniques can also provide a precise view of the vascularisation of the hippocampal region, but morphometric studies are difficult and the lack of visualization of the cerebral parenchyma makes interpretation difficult with this technique [6]. Segmentation of HiP-CT images allowed for quantitative vascular measurements from the brain structure. Synchrotron HiP-CT has underscored its value in micro-anatomical studies of cerebral and vascular structures, eliminating the need for contrast agents and avoiding tissue damage. This technique has already been used to study the entire arterial network of an organ such as the kidney [19], and has been used to visualise thrombosed microvessels in lungs infected with SARS-CoV-2 [1].

We successfully visualised the intricate structure of the hippocampus arterial tree on two brains, including intraparenchymal arteries, and constructed a 3D representation of this vascular network. This advanced imaging method enabled the identification of vessels as small as 7 μm in diameter. In accordance with the literature, our observations showed vascularisation predominantly originating from the PCA [4] with multiple anastomoses in the hippocampal arterial network, augmenting resistance to anoxia. We did not identify any hippocampal branches arising from the anterior choroidal artery, possibly due to the variability in hippocampal vascularisation [4, 13] and our analysis being limited to two brains. The anterior choroidal artery appears to play a secondary role in the vascularisation of the hippocampus, although this artery and its perforating branches pose a risk of cerebral infarction during hippocampal resection following temporal lobe epilepsy [6]. Regarding the hippocampal arteries themselves, based on our observations and in line with recent literature, it is more accurate to refer to arterial complexes formed by a dense anastomotic network rather than three distinct hippocampal arteries [26]. However, our observations did not reveal the vascular arcade in the hippocampal sulcus, which seems to be present in 23% of subjects [26]. Additionally, we did not find evidence of hippocampal devascularisation originating from the posterior choroidal arteries, as is sometimes described [10].

Limitations of this work include the limited throughput of the acquisition system, the resolution boundary of the organ-wide scan, and the current restricted accessibility to the technique. Regarding the limitations of our imaging findings, it should be noted that our observations pertain to only two brains, focusing on an anatomical structure known for its high variability. As such, our results should not be considered generalisable. However, this study represents the inaugural attempt to image the human hippocampus at such a high-resolution using synchrotron techniques. We successfully identified and traced the superficial hippocampal artery, though we were limited in tracking the intraparenchymal vessels comprehensively. The post-mortem nature of our specimen, along with the pre-casting treatments prior to imaging, might have introduced alterations to the native brain and vascular structures. Damage of the brain was noted following the vacuum degassing procedure used for the LADAF-2020-31 brain, although this did not affect the hippocampus. The LADAF-2021-17 brain was degassed using thermal cycling instead to avoid such damage [2]. Additionally, the segmentation performed in this work was largely manual as it could not be fully automated. This is due to the intricacy of the structures involved and the presence of numerous collapsed arterioles; future advancements might consider integrating deep learning segmentation algorithms [27] as well as injection of resin in the vascular network at the beginning of the sample preparation to avoid the collapse of large vessels. While this research emphasised the arterial network of the hippocampus, venous networks offer another potential avenue of exploration in future work.

## Conclusion

Synchrotron HiP-CT has emerged as a key non-destructive imaging method offering high-resolution visuals of large structures, preserving anatomical relationships, in a three-dimensional perspective. This enabled us to discern intraparenchymal hippocampal arteries and construct a three-dimensional representation of them. This study has highlighted the efficiency of HiP-CT for anatomical exploration on two human brains, specifically vascular study, down to arteries in the few micrometres range. Moreover, the HiP-CT is also intended in the future to be used to improve in-vivo medical imaging by using synchrotron data as a reference to train super-resolution algorithms to be integrated into conventional machines. Such perspectives could increase our micro-anatomical knowledge, help guide surgery and improve our understanding of hippocampal pathologies.

## Acknowledgments

The authors sincerely thank those who donated their bodies to science so that anatomical research could be performed. Results from such research can potentially increase humanity’s overall knowledge that can then improve patient care. Therefore, these donors and their families deserve our highest gratitude.

The authors would like to express their gratitude for the financial support provided by the Chan Zuckerberg Initiative (2020-225394, CZIF2021-006424), an advised fund of SVCF, the MRC (MR/R025673/1), and ESRF beamtimes (md1252 & md1290). P.D.L. is supported by a Royal Academy of Engineering Chair in Emerging Technologies (CiET1819/10).

## Data sharing statement

The first complete brain scan at 25.08um was already made public with the initial HiP-CT publication in 2021 in the Human Organ Atlas (https://human-organ-atlas.esrf.eu): DOI 10.15151/ESRF-DC-572252655. The additional zooms have been available in the Human Organ Atlas with the respective DOIs 10.15151/ESRF-DC-572253460 for the 6.05um scan, and 10.15151/ESRF-DC-572252431 for the 2.45um scan. The second complete brain was not yet available to the public. The scans of this second brain will be released on the Human Organ Atlas with publication of the present paper.

## Compliance with Ethical Standards

As part of research involving human participants in the context of body donation, the authors declare that they have complied with current French law (funeral and public health legislation) and have respected the ethical principles of respect for the deceased. The research was conducted in accordance with the 1964 Helsinki Declaration. The research was authorised by the Grenoble Alpes University.

## Statement about authors’ contribution

Alexandre BELLIER: protocol/project development, sample acquisition and preparation, data collection or management, data analysis, manuscript writing/editing

Paul TAFFOREAU: protocol/project development, imaging technique development, sample preparation, data collection or management, data analysis, manuscript writing/editing

Ali BOUZIANE: data analysis, manuscript writing/editing

Tanguy ANGELLOZ-NICOUD: data analysis, manuscript writing/editing

Peter D. LEE: protocol/project development, data collection or management, manuscript writing/editing

Claire WALSH: protocol/project development, data collection or management, manuscript writing/editing

